# Oral cholestyramine prevents enrichment of diverse daptomycin-resistance mutations in intestinal *Enterococcus faecium*

**DOI:** 10.1101/2022.05.25.493495

**Authors:** Valerie J. Morley, Derek G. Sim, Aline Penkevich, Robert J. Woods, Andrew F. Read

**Affiliations:** Center for Infectious Disease Dynamics, Department of Biology, The Pennsylvania State University, University Park, PA, USA; Center for Global Health, Department of Internal Medicine, The University of New Mexico Health Sciences Center, Albuquerque, NM, USA; Division of Infectious Diseases, Department of Internal Medicine, University of Michigan, Ann Arbor, MI, USA; Department of Entomology, The Pennsylvania State University, University Park, PA, USA; Huck Institutes for the Life Sciences, The Pennsylvania State University, University Park, PA, USA

## Abstract

**Background and Objectives:** Previously, we showed proof-of-concept in a mouse model that oral administration of cholestyramine prevented enrichment of daptomycin-resistant *Enterococcus faecium* in the gastrointestinal (GI) tract during daptomycin therapy. Cholestyramine binds daptomycin in the gut, which removes daptomycin selection pressure and so prevents the enrichment of resistant clones. Here, we investigated two open questions related to this approach: 1) can cholestyramine prevent the enrichment of diverse daptomycin mutations emerging *de novo* in the gut? 2) how does the timing of cholestyramine administration impact its ability to suppress resistance?

**Methodology:** Mice with GI *E. faecium* were treated with daptomycin with or without cholestyramine, and *E. faecium* was cultured from feces to measure changes in daptomycin susceptibility. A subset of clones was sequenced to investigate the genomic basis of daptomycin resistance.

**Results:** Cholestyramine prevented the enrichment of diverse resistance mutations that emerged *de novo* in daptomycin-treated mice. Whole-genome sequencing revealed that resistance emerged through multiple genetic pathways, with most candidate resistance mutations observed in the *clsA* gene. Additionally, we observed that cholestyramine was most effective when administration started prior to the first dose of daptomycin. However, beginning cholestyramine after the first daptomycin dose reduced the frequency of resistant *E. faecium* compared to not using cholestyramine at all.

**Conclusions and Implications:** Cholestyramine prevented the enrichment of diverse daptomycin-resistance mutations in intestinal *E. faecium* populations during daptomycin treatment, and it is a promising tool for managing transmission of daptomycin-resistant *E. faecium*.

## Lay Summary

We demonstrated experimentally in a mouse model that administration of the FDA-approved drug cholestyramine prevented the spread of daptomycin resistance in bacterial populations in the gut. This is important because resistant bacteria in the gut can be transmitted to other individuals or be a source of infections. It might be possible to use cholestyramine as an adjuvant to prevent the emergence of daptomycin-resistant bacteria in patients being treated intravenously with daptomycin.

## Background and Objectives

As the antimicrobial resistance threat grows, it is critical to find ways to use antimicrobials in healthcare without fueling the spread of resistance. With this goal in mind, we recently showed proof-of-concept for a strategy to reduce the transmission of daptomycin-resistant *Enterococcus faecium* during daptomycin therapy [1]. Daptomycin is administered intravenously to treat blood and soft tissue infections and is one of the few antibiotics indicated for the treatment of vancomycin resistant *E. faecium* (VRE) [2]. During daptomycin therapy, most daptomycin is renally excreted, but 5-10% of the dose enters the intestines [3]. This means *E. faecium* colonizing the intestines are exposed to daptomycin, which can select for resistance in intestinal populations [1,4]. This off-target daptomycin selection could be an important source of hospital-transmitted daptomycin-resistant VRE because *E. faecium* transmission is fecal-oral [5]. We hypothesized that inactivating daptomycin locally in the intestines could prevent the enrichment and transmission of daptomycin-resistant *E. faecium*, while still allowing daptomycin therapy to effectively target infections in sites like the bloodstream. We tested the potential of the FDA-approved drug cholestyramine to be repurposed as a daptomycin antagonist to inactivate daptomycin in the gut. In a previous paper, we used a mouse model to show that pairing oral cholestyramine with systemic daptomycin treatment dramatically reduced the fecal shedding of daptomycin-resistant *E. faecium* [1]. Cholestyramine treatment is therefore a promising strategy that allows for daptomycin therapy while blocking the enrichment and fecal shedding of daptomycin-resistant *E. faecium*.

Our previous paper provided proof-of-concept that cholestyramine adjuvant therapy could limit the spread of daptomycin resistance. That work left two major open questions that we address here in the same mouse model. First, can cholestyramine prevent the enrichment of resistance mutations that arise *de novo* in the gut during treatment? Our previous paper showed that cholestyramine successfully prevented the enrichment of daptomycin-resistant clones that were experimentally inoculated at low levels in intestinal populations. Preventing *de novo* resistance poses additional challenges, as there may be larger mutational inputs from larger sensitive populations, and a greater diversity of resistance genotypes and phenotypes may need to be suppressed. Second, how does the timing of cholestyramine administration affect resistance emergence? In our previous experiments, we began administering cholestyramine to mice one day prior to their first daptomycin injection. In clinical settings, it might not be practical to start cholestyramine treatment prior to the first dose of daptomycin.

## Methodology

### Experimental overview

We conducted two experiments. The first experiment tested the ability of cholestyramine to prevent the enrichment of resistance mutations that occurred *de novo* during treatment. We treated 180 mice colonized with daptomycin-susceptible *E. faecium* with either 50 or 100 mg/kg subcutaneous daptomycin daily for 5 days. Half the mice were provided with a diet supplemented with cholestyramine starting one day prior to the first daptomycin injection; the other half were provided the same diet but without cholestyramine. We sequenced a subset of the resistant clones that emerged to determine likely resistance mechanisms.

The second experiment tested whether cholestyramine effectively inhibited the enrichment of resistance if cholestyramine treatment and daptomycin treatment began at the same time. Groups of ten mice were inoculated orally with a mixture of daptomycin-susceptible and daptomycin-resistant *E. faecium* (95% susceptible, 5% resistant). This design drastically decreases the number of mice needed compared to measuring *de novo* resistance emergence because it eliminates the wait time for resistance mutations in the *E. faecium* population and focuses on enrichment for the resistant clone under daptomycin selection pressure. Mice were treated with 50 mg/kg daptomycin for 5 days. To test the effect of cholestyramine therapy timing, mice received either no cholestyramine, a cholestyramine-supplemented diet starting one day prior to daptomycin treatment (‘early’), or a cholestyramine-supplemented diet starting shortly after their first daptomycin dose (‘late’).

In both experiments, the enrichment of resistance was the key readout. As in our earlier paper [1], this was measured as the percent of *E. faecium* in fecal samples that were daptomycin resistant, and the total density of resistant *E. faecium* in feces (number per gram).

### Mice and bacterial strains

Mice in all experiments were female Swiss Webster (CFW) obtained from Charles River Labs. This is an outbred mouse line. Mice were fed a standard diet (5001 Laboratory Rodent Diet) or a standard diet supplemented with 2% w/w cholestyramine resin (Sigma-Aldrich catalog #C4650). To minimize bacterial cross-contamination between animals, all mice were housed individually during experiments, and experimenters changed gloves between handling different mice.

We used the daptomycin-susceptible *E. faecium* strain BL00239-1 (MIC_c_ = 3.1 (Minimum Inhibitory Concentration computed, see below)), a vancomycin resistant strain originally isolated from a bloodstream infection at the University of Michigan Hospital. In the experiment testing the timing of cholestyramine administration, we also used the daptomycin-resistant strain BL00239-1-R (MIC_c_ = 8.6). This resistant strain was isolated from a mouse colonized with BL00239-1 after experimental daptomycin treatment, as previously described [1].

### Experimental detail

Animal experiments were approved by the Institutional Animal Care and Use Committee at the Pennsylvania State University. All mice were pretreated with ampicillin (0.5 g/L in drinking water) for 7 days before *E. faecium* inoculation. Ampicillin disrupts the natural gut microbiota and facilitates *Enterococcus* colonization [6]. Mice that were co-housed during ampicillin pre-treatment were evenly allocated among experimental treatment groups. *E. faecium* strains were plated from glycerol stocks and then grown overnight in liquid culture in Brain Heart Infusion broth. Mice were inoculated via oral gavage with 10^8^ CFU *E. faecium* suspended in saline. *E. faecium* inoculum counts were confirmed by plating. Following *E. faecium* inoculation, mice were split into individual cages with untreated water and experimental diets. Daptomycin doses were administered daily starting one day post inoculation via subcutaneous injection. Daptomycin doses were based on an average mouse weight for each experiment. For mice receiving cholestyramine, a cholestyramine supplemented diet (2% w/w) was provided to mice starting one day prior to the first daptomycin dose or starting the same day as the first daptomycin dose, as described in experimental results. Mice were allowed to eat *ad libitum*, and once the cholestyramine was provided, the mice continued to be maintained on this diet for the duration of the experiment. For stool collection, mice were placed in clean plastic cups, and fresh stool was collected using a sterile toothpick. Stool samples were suspended in PBS (25 uL PBS/mg stool) and frozen with glycerol at -80°C for subsequent analysis.

### Analysis of E. faecium in stool samples

*E. faecium* were enumerated by plating diluted fecal suspensions on selective plates (Enterococcosel agar supplemented with 16 ug/mL vancomycin). Plates were incubated at 35°C for 40-48 hours, and colonies were counted. To quantify the proportion of these bacteria that were daptomycin resistant, fecal suspensions were plated on calcium-supplemented Enterococcosel plates with 16 ug/mL vancomycin and 10 ug/mL daptomycin. Plates were incubated at 35°C for 40-48 hours, and colonies were counted. Serially-diluted fecal suspensions were each plated once on plates without daptomycin and once on plates containing daptomycin to estimate the proportion of daptomycin-resistant bacteria. The limit of detection was 20 CFU per 10 mg feces.

In experiment 1, a subset of *E. faecium* clones were isolated from fecal samples and analyzed by broth microdilution. Clones were purified by streaking twice on Enterococcosel agar with 16 ug/mL vancomycin and were then stored in glycerol stocks at -80°C. Broth microdilutions were performed according to Clinical & Laboratory Standards Institute (CLSI) guidelines [7]. After incubation, cell densities were measured by OD600 absorbance in a plate reader. OD values were fitted to a Hill function to determine the computed MIC (MIC_c_), the concentration at which the Hill function crossed a cutoff two standard deviations above the mean of the negative control wells, as described previously [4].

### Genome sequencing of E. faecium isolates

In experiment 1, daptomycin resistance was observed in >5% of the *E. faecium* population in 22 of 180 mice using culture-based methods. We isolated 5 clones from each of these 22 mice for susceptibility testing. From this set of isolates, 4 clones from each mouse were selected for whole-genome sequencing (89 total isolates sequenced including the ancestral BL00239-1 clone). The clone selection process for sequencing was as follows: from each mouse we chose the clones with the highest and lowest MIC_c_s, and then two additional clones were selected randomly from each mouse using a random number generator.

Sequencing libraries were prepared from whole genomic DNA using the Collibri PCR-free ES DNA Library Prep Kit for Illumina with UD Indexes (sets A-D 1-96). Libraries were submitted to the University of Michigan sequencing core for paired-end 150bp sequencing on the Illumina NovaSeq 6000. Sequencing reads were trimmed using Trimmomatic [8], and MultiQC was used to assess data quality [9]. One isolate (M3.5-5) was excluded from downstream analysis due to poor sequence quality. Trimmed reads were mapped against the BL00239-1 reference genome (generated previously [1]) and the *E*. faecium DO reference genome (NC_017960.1) using Burrows-Wheeler Aligner (BWA) [10], and candidate variants were identified with The Genome Analysis Toolkit (GATK) [11]. Candidate variants were annotated using SnpEff [12]. Reads from the ancestral clone were aligned to the reference genome (aligned to self or DO) to generate a list of background variants; these background variants were filtered out during variant calling. Remaining candidate variants were screened by visual inspection of alignments in Tablet [13].

### Statistical analysis

Statistical analyses were run in R v1.2.1335 [14] using the packages ‘nlme’ [15] and ‘glmmTMB’ [16]. To analyze proportions of resistant bacteria, samples were plated on agar with and without daptomycin, resulting in a count of resistant bacteria and a count of total bacteria. Due to sampling, these proportions were not bounded by one, so proportion data were normalized by dividing each value by the maximum value in the data set. Proportions of resistant bacteria were analyzed using mixed binomial regression models. Absolute densities of vancomycin-resistant *E. faecium* were analyzed using mixed models with an autoregressive error structure as previously described [17]. In experiment 2, total number of bacteria or resistant bacteria shed through time were estimated from the area under the density through time curves. Full model structures and output are shown in Supplementary File 1. Figures were created with ggplot2 [18].

## Results

### Cholestyramine prevented the enrichment of diverse daptomycin resistance mutations

Fecal samples were collected at Day 1 (prior to daptomycin) and Day 8 (after daptomycin treatment), and the fecal densities of total *E. faecium* and daptomycin-resistant *E. faecium* were determined (Supplementary Figure 1). For mice with no detectable *E. faecium* in the Day 8 fecal sample (n=19), an additional sample was collected at Day 14.

**Fig 1.**
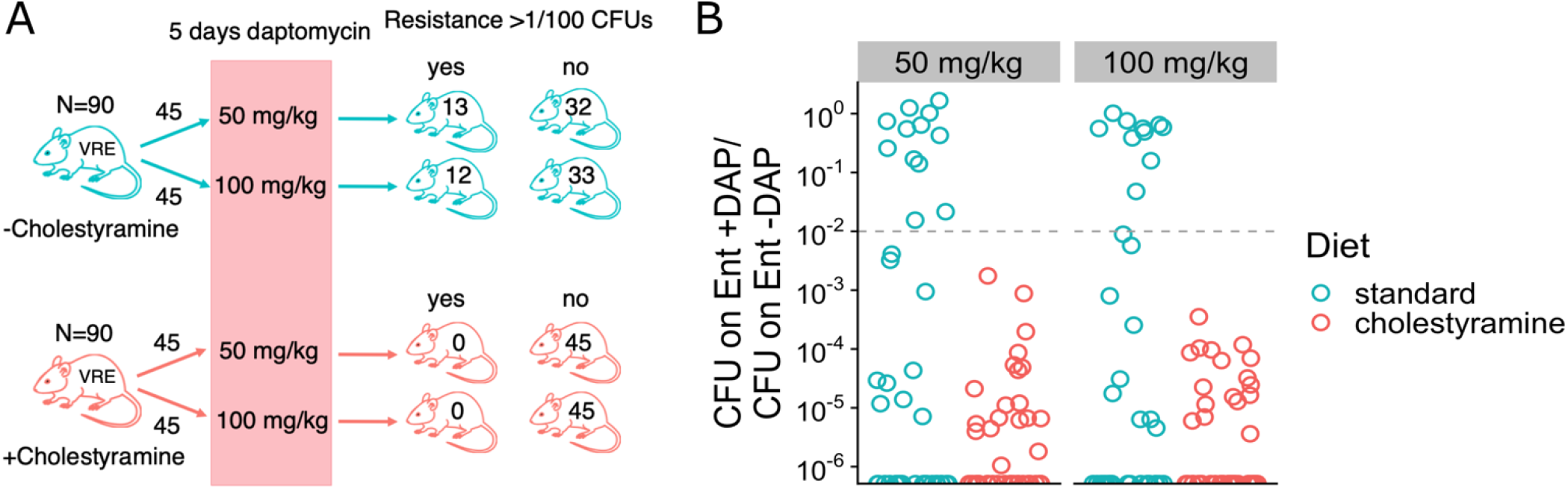
Cholestyramine inhibits enrichment of daptomycin resistance mutations in gut *E. faecium* populations. A) Experimental design. Mice colonized with daptomycin-susceptible *E. faecium* were supplied with a standard diet or a diet supplemented with 2% cholestyramine by weight (N = 90 per diet treatment). Mice then received subcutaneous daptomycin for 5 days at 50 or 100 mg/kg daily. Fecal samples were collected at Day 8, and the frequency of daptomycin-resistant *E. faecium* in these fecal samples was determined by culture. For mice with low *E*. faecium shedding at Day 8 (n=19), an additional sample was collected at Day 14. Numbers show how many mice had >1/100 daptomycin-resistant *E. faecium* in a Day 8 or Day 14 fecal sample. B) Proportion of daptomycin-resistant *E. faecium* at Day 8. Fecal samples were plated on Enterococcal agar plates (Ent -DAP) and on Enterococcal plates supplemented with daptomycin (Ent +DAP). The proportion of daptomycin-resistant *E. faecium* was estimated as (CFU on Ent +DAP)/(CFU on Ent -DAP). Each point represents a fecal sample from one mouse. The dotted line shows 1/100 CFUs resistant, as in Figure 1A.

Daptomycin-resistant mutants were detected in 48% (86/180) of all mice and were detected in similar numbers or mice in both groups (cholestyramine: 37/90, no cholestyramine, 49/90; Pearson’s chi-square test p = 0.10). However, the frequency of daptomycin resistance in the *E. faecium* population within a mouse never rose above 1/100 CFUs in any of the 90 mice treated with cholestyramine. In contrast, it rose above 1/100 frequency in 28% (25/90) of mice treated with daptomycin in the absence of cholestyramine, and often came to dominate (Figure 1). Resistance was significantly less likely to rise above 1/100 in cholestyramine-treated mice (Pearson’s chi-square test p < 0.001). Thus, cholestyramine successfully prevented *de novo* emergence of daptomycin resistance in gastrointestinal (GI) *E. faecium* populations.

### Cholestyramine prevented enrichment of resistance that emerged through multiple genetic pathways

To confirm the emergence of resistance, we determined the daptomycin MIC_c_ for a subset of *E. faecium* isolates. For the 22 mice which had resistance 5/100 CFUs at Day 8 or Day 14, we isolated 5 *E. faecium* clones from each mouse’s fecal sample and estimated MIC_c_ by broth microdilution. The broth microdilution assay confirmed that at least 16 mice harbored *E. faecium* isolates with elevated daptomycin MIC_c_ relative to the ancestral clone (Fig 2).

**Fig 2.**
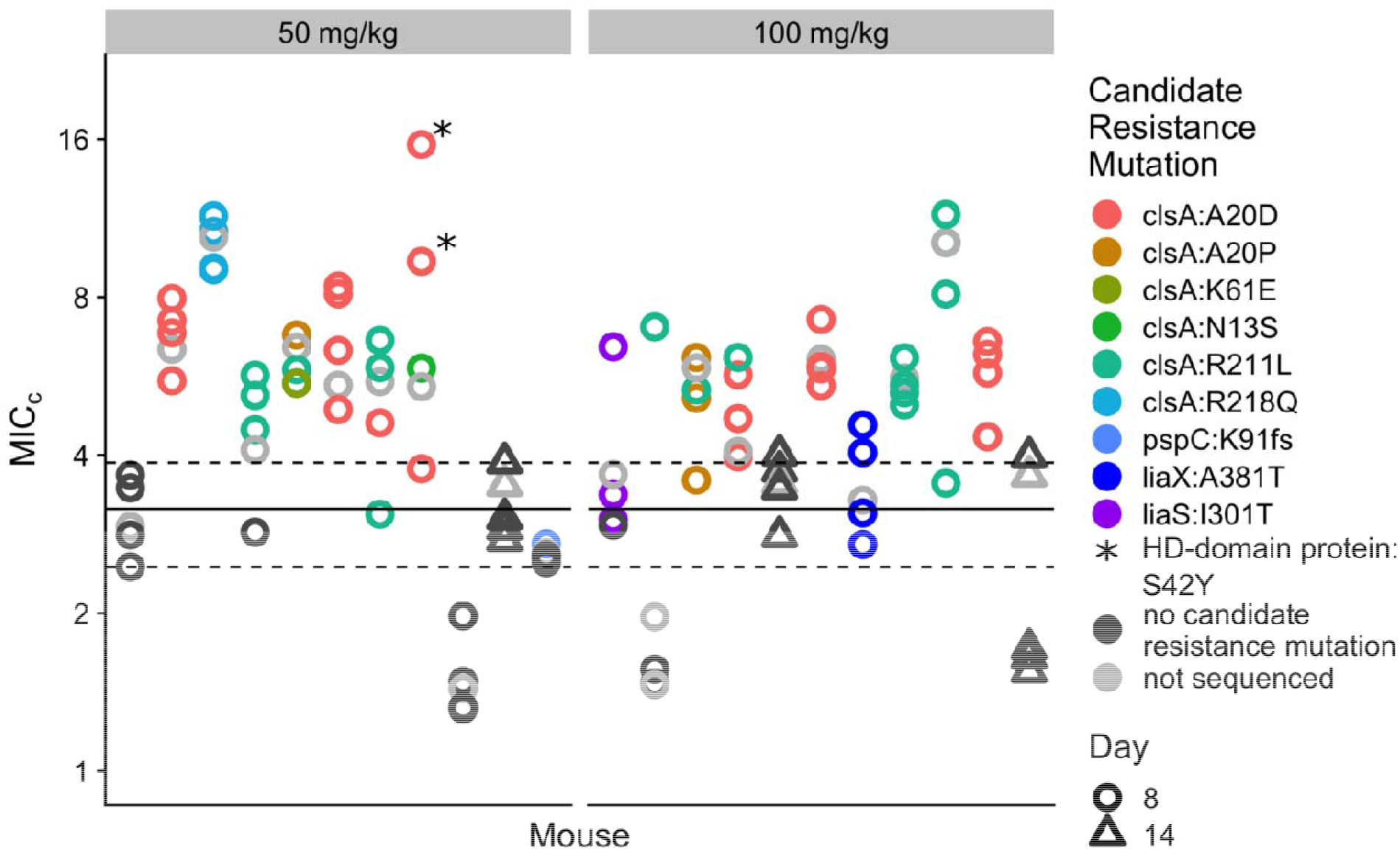
Daptomycin resistance in *E. faecium* clones isolated from mouse fecal samples. 22 mice were identified as having *E. faecium* populations with increased daptomycin resistance based on an initial plating assay (>5/100 CFUs resistant at Day 8 or Day 14). 5 *E. faecium* clones were isolated from each of these fecal samples. Daptomycin MIC_c_s were determined by broth microdilution for each clone (see methods). Each point shows the MIC_c_ of a single clone (mean of 2 measurements). The line shows the MIC_c_ of the ancestral clone BL00239-1 (mean of 4 measurements) with 95% CI (dotted lines). Daptomycin doses are indicated at the top of each panel. A subset of these isolates was sequenced, and colors indicate mutations in genes associated with daptomycin resistance. The isolates marked with an asterisk had 2 candidate resistance mutations in *clsA* and an HD-domain containing protein.

To investigate the genetic mechanism of resistance in these isolates, we conducted whole-genome sequencing on 4 clones from each of these 22 mice (89 isolates total including the ancestral clone). A full list of observed mutations can be found in Supplementary Table 1. Across the sequenced isolates, we found mutations in several genes known to be involved in daptomycin resistance (Fig 2). The most common candidate resistance mutations were found in the *clsA* gene, which has previously been associated with daptomycin resistance in *E. faecium* [19–23]. The *clsA* gene encodes cardiolipin synthase, which produces cardiolipin. Increased cardiolipin content in the bacterial membrane inhibits membrane permeabilization by daptomycin [19]. We found *clsA* mutations in isolates from 14 of the 22 mice. Six different *clsA* mutations were observed, all of which were nonsynonymous. The most common *clsA* mutation was A20D, followed by R211L. Within 4 of the 14 mice with *clsA* mutations, we observed two with different mutations in different clones from the same fecal sample, and in one mouse we observed three different *clsA* mutations. This means that multiple *clsA* mutations appeared independently within these populations.

In 2 mice, we also observed mutations in *liaS* (I301T) and *liaX* (A381T), another set of genes associated with daptomycin resistance [21,23,24]. The three-component LiaFSR system regulates the cellular envelope stress response, which impacts daptomycin susceptibility [25]. The three-gene cluster *liaXYZ* is hypothesized to be regulated by *liaR* [24]. Additionally, we observed mutations in an HD-domain containing protein previously associated with daptomycin resistance (S42Y) [21] and frameshift in the candidate resistance gene *pspC* [21].

In summary, daptomycin resistance emerged through multiple genetic mechanisms, with strong convergence at the gene level (*clsA*). Cholestyramine prevented the enrichment of resistance in all treated mice, despite the availability of multiple genetic pathways to resistance in this system.

### Early cholestyramine administration better prevented the enrichment of resistance

Both early and late cholestyramine treatments significantly reduced the proportion of *E. faecium* that were daptomycin-resistant relative to no-cholestyramine controls (Fig 3A; mixed binomial model, early*Day p=0.006, late*Day p=0.03, Model 1 in Supplementary File 1). ‘Early’ cholestyramine had a substantially larger effect size at all time points (Fig 3A, Model 1 in Supplementary File 1). Only the ‘early’ treatment significantly reduced total shedding of daptomycin-resistant *E. faecium*, as measured by area under the curve of density through time (AUC) (Fig 3B; generalized linear model (glm), early p=0.01, late p=0.12, Model 2 in Supplementary File 1). ‘Early’ cholestyramine reduced absolute shedding of resistant *E. faecium* (AUC) by 92% compared to no cholestyramine.

**Fig 3.**
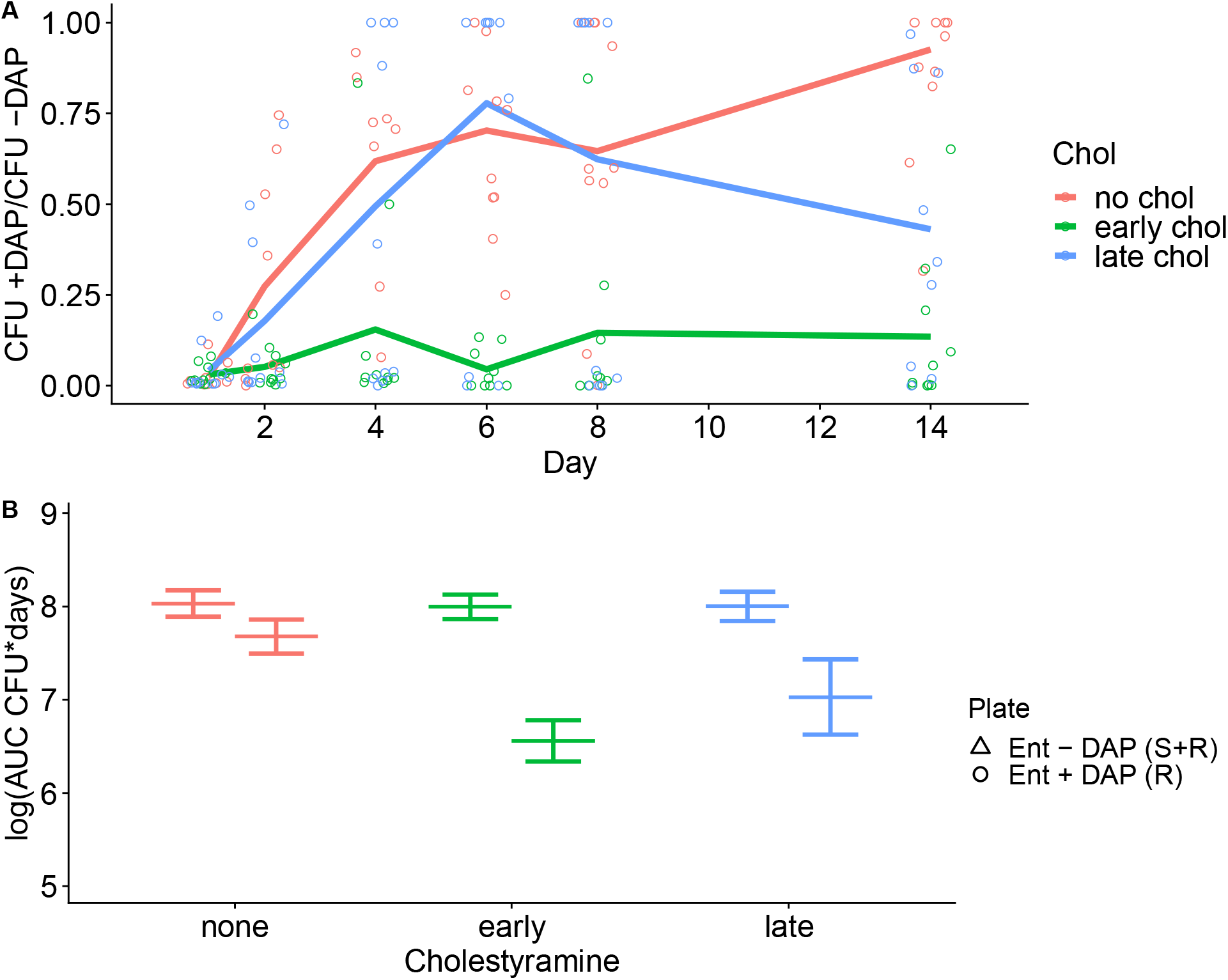
Effect of timing of cholestyramine administration on enrichment for resistance. Mice treated with daptomycin were provided with a standard diet (no chol), cholestyramine starting one day prior to the first daptomycin injection (early chol), or cholestyramine starting immediately after the first daptomycin injection (late chol) (N=10 per treatment). Fecal samples were collected at time points during and after daptomycin treatment, and densities of *E. faecium* and daptomycin-resistant *E. faecium* were determined by plating. A) The proportion of *E. faecium* in mouse fecal samples that were daptomycin-resistant (CFU +DAP/CFU -DAP). Means are shown, with points indicating values for individual samples. Note that fecal samples with no detectable *E. faecium* (n=16) were excluded from this analysis. B) The area under the curve of density through time (AUC) for the absolute densities of total *E. faecium* and daptomycin-resistant *E. faecium* in fecal samples over the total duration of the experiment. Points indicate AUCs for individual mice for total *E. faecium* (triangles, measured on Enterococcosel plates containing daptomycin) and daptomycin-resistant *E. faecium* (circles, measured on Enterococcosel plates without daptomycin). Means and SE are shown (N=10).

In daptomycin-treated mice, cholestyramine treatment did not affect total shedding of *E. faecium* (Fig 3B; glm, early p=0.86, late p=0.88, Model 3 in Supplementary File 1). In control mice that received no daptomycin but received identical cholestyramine treatments, cholestyramine alone also did not affect total *E. faecium* shedding (Supplementary Figure 2; glm, early p=0.19, late p=0.22, Model 4 in Supplementary File 1).

## Conclusions and Implications

Here, we have shown that cholestyramine can prevent enrichment of resistance following the *de novo* emergence of daptomycin-resistance in *E. faecium* populations colonizing the intestines (Experiment 1, Fig 1). Previously, we had shown that cholestyramine could prevent the enrichment of preexisting resistant clones [1] (a result we saw again in Experiment 2; Figure 3). These results further show the promise of cholestyramine treatment for preventing off-target resistance evolution emerging via multiple genetic pathways during daptomycin treatment. Ideally, cholestyramine would allow clinicians to effectively treat bloodstream and soft tissue infections with daptomycin, while preventing the emergence of daptomycin-resistant *E. faecium* from intestinal populations and risk of onward transmission.

Daptomycin resistance in *E. faecium* arises through chromosomal mutations and there are many possible genetic pathways to increased resistance [21]. This means that within a patient, especially in large *E. faecium* populations colonizing the gut, there is a high probability that daptomycin resistance mutations will appear through random mutation. Treatment then imposes selection, which can enrich for these resistant clones. This process of resistance emergence has been observed in hospital patients—*E. faecium* isolated from patient perirectal swabs are more resistant following daptomycin treatment, and genomic data suggests that this resistance arises *de novo* within patients [4]. Here, we observe the same process in a mouse model. We detected the presence of *de novo* resistance (at least one resistant clone) in 48% of mice (Figure 1). In the absence of cholestyramine, daptomycin treatment resulted in the up-selection of this resistance above 1/100 CFUs in about half of those mice (Fig 1). Resistance emerged so frequently likely because mutations can confer resistance via multiple genetic pathways (Fig 2). Should similar rates of spontaneous resistance mutation and emergence occur in patients colonized with *E. faecium*, adjuvant therapies that can inhibit daptomycin activity in the GI tract could have substantial impact. Importantly, cholestyramine effectively suppressed resistance emergence even though resistance was readily accessible through multiple genetic pathways (Figs 1, 2). We hypothesize that cholestyramine works by eliminating selection pressure for daptomycin resistance, thus preventing the enrichment of resistant clones, independent of the genetic mechanism of resistance.

These results also show that the timing of cholestyramine administration may matter. We found in mice that cholestyramine was most effective at preventing resistance when administration began 24 hours before the first daptomycin dose, although beginning cholestyramine after the first daptomycin dose still had some suppressive effect on the frequency of resistance. An essential step in translating cholestyramine treatment to human subjects will be understanding the pharmacokinetics of cholestyramine and daptomycin, especially when the drugs reach the intestines after administration and in what quantities. This pharmacokinetic data should inform decisions about the optimal cholestyramine regimens.

Cholestyramine is one of several adjunctive therapies in development that could prevent off-target resistance emergence in the gut during antibiotic treatment [26–29]. These treatments rely on antibiotic antagonists, which bind to or inactivate antibiotics locally in the intestines. Daptomycin is an especially attractive antibiotic target, because daptomycin resistance emerges easily and often in the gut through point mutations, as shown here in our mouse model and in patient data [4]. Additionally, cholestyramine is an attractive drug to repurpose as an adjuvant, because it is an FDA-approved drug that has been in use with minimal side effects for over 50 years [30,31], and it could inexpensively be repurposed to prevent the spread of resistance. These adjunctive therapies are exciting new tools for antimicrobial stewardship, which offer the possibility of using antibiotics to treat patients without the onward transmission of drug-resistant pathogens.

## Supporting information

Supplemental Figues

Supplementary File 1 - Statistical Models

## Acknowledgements

We thank Kevin Tracy for assistance in generating the nanopore sequencing and analysis pipeline.

## Data availability

Data are available on Dryad (DOI https://doi.org/10.5061/dryad.rxwdbrvbm). Raw sequencing reads are available through NCBI Bioproject PRJNA832698.

