## Supplemental Figues for "Oral cholestyramine prevents enrichment of diverse daptomycin-resistance mutations in intestinal *Enterococcus faecium*"

**Supplementary Figures:**

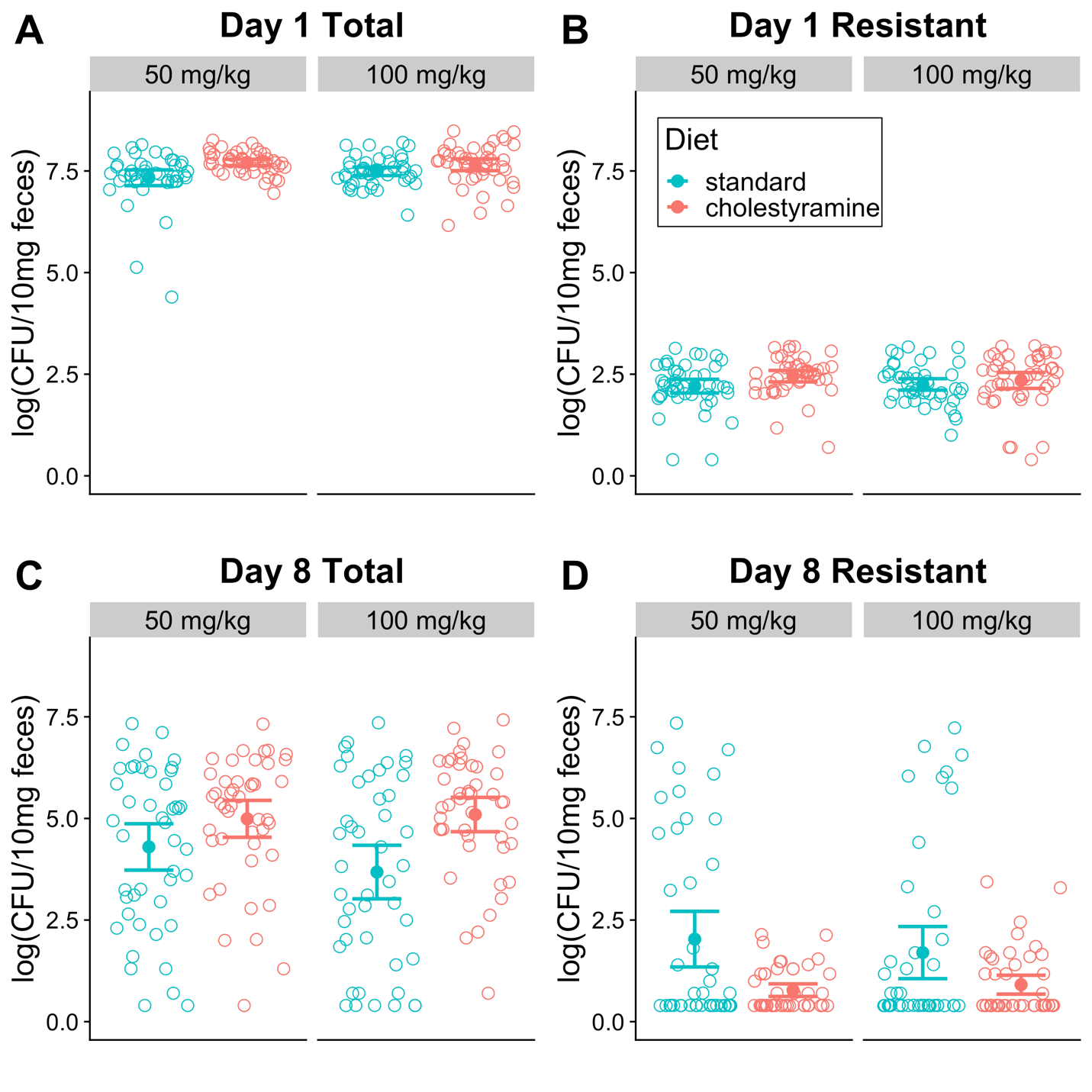

**Supp Fig 1.** **Densities of *E. faecium* in mouse feces during *de novo* emergence experiment.** Each point shows the density of *E. faecium* in a fecal sample from one mouse. Densities were determined by plating on Enterococcosel agar with and without daptomycin supplementation. Mean and 95% CI shown. Samples were analyzed at Day 1 (prior to daptomycin treatment) and Day 8 (post daptomycin treatment) A) Total *E. faecium* densities at Day 1. B) Densities of daptomycin-resistant *E. faecium* at Day 1. C) Total *E. faecium* densities at Day 8. D) Densities of daptomycin-resistant *E. faecium* at Day 8.

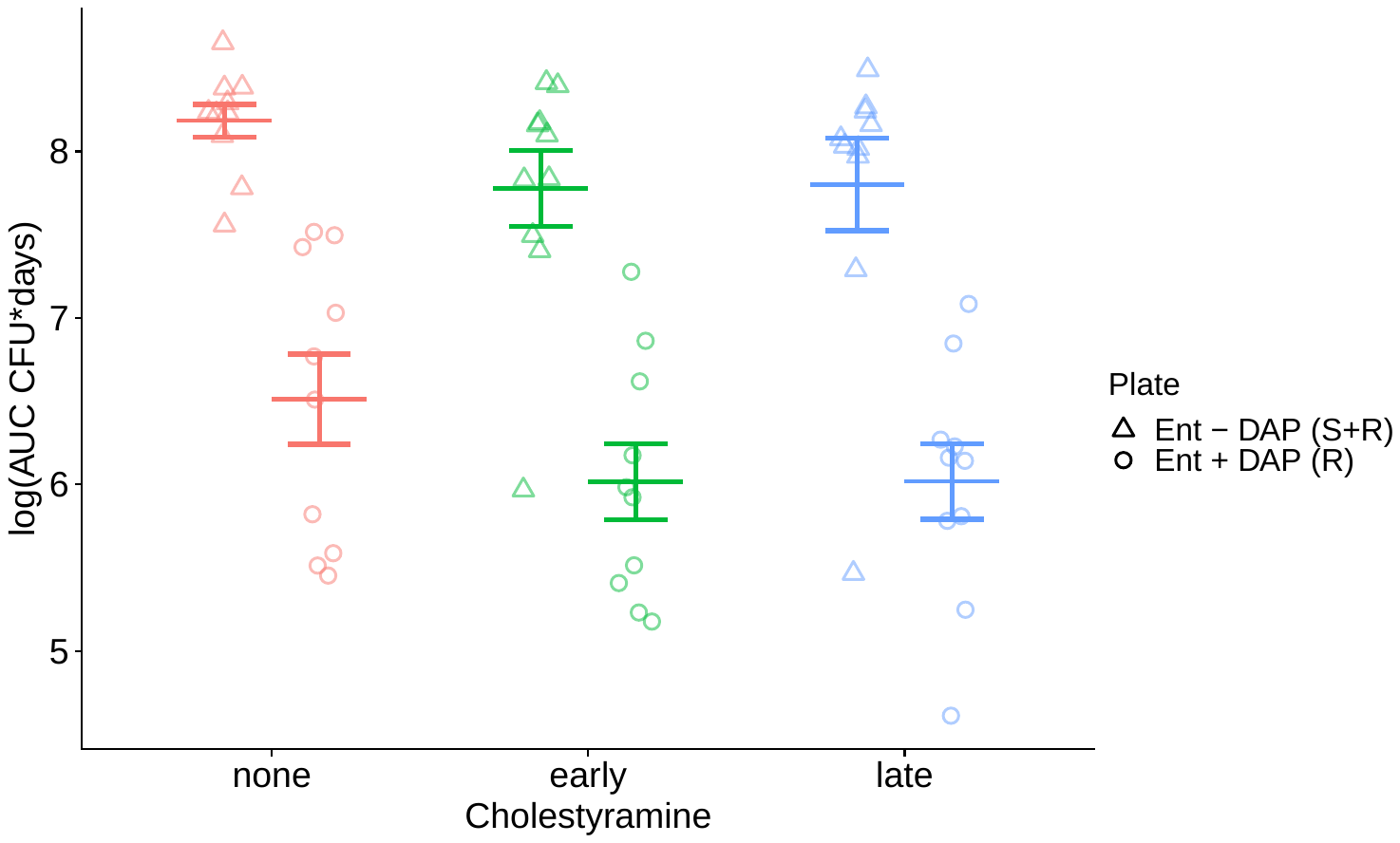

**Supp Fig 2.** **Effect of cholestyramine alone on *E. faecium* densities in experiment 2.** As part of the experiment testing the timing of cholestyramine administration, control mice were provided with cholestyramine in the absence of daptomycin treatment. Mice were provided with a standard diet (no chol), cholestyramine starting at Day 0, or cholestyramine starting at Day 1 (late chol) (N=10 per treatment). Fecal samples were collected at time points throughout the experiment, and densities of *E. faecium* and daptomycin-resistant *E. faecium* were determined by plating. Plot shows the area under the curve (AUC) for the absolute densities of total *E. faecium* and daptomycin-resistant *E. faecium* in fecal samples over the total duration of the experiment. Points indicate AUCs for individual mice for total *E. faecium* (triangles, measured on Enterococcosel plates containing daptomycin) and daptomycin-resistant *E. faecium* (circles, measured on Enterococcosel plates without daptomycin). Means and SE are shown (N=10).
