## Supplementary File 1 - Statistical Models for "Oral cholestyramine prevents enrichment of diverse daptomycin-resistance mutations in intestinal *Enterococcus faecium*"

**Supplementary File 1 - Statistical Models for Experiment 2**

**Model 1: Effect of cholestyramine timing on proportion of daptomycin-resistant *E. faecium***

**Input data:**Fig3_CFUdata_choltiming.csv Dryad DOI <https://doi.org/10.5061/dryad.rxwdbrvbm>

**Data preparation:**

1. Coded variable Day as factor (Day.factor)
2. Created a subset of the data (data.sub) that omits entries where total.per10mg = 0
3. Created variable for proportion resistant (Prop). This is resist.per10mg/ total.per10mg. This variable was normalized by dividing by the maximum value of Prop to create the normalized variable (PropNorm).

**Model Structure in R:**

glmmTMB(PropNorm~Chol*Day + (1|Mouse_ID) + ar1(Day.factor + 0 | Mouse_ID),

data = na.omit(data.sub[data.sub$Day>1&data.sub$Antibiotic=="daptomycin",]),

family = binomial, weights = total.per10mg,

control = glmmTMBControl(optimizer = optim, optArgs = list(method="BFGS")))

summary()

|  | Estimate | Std. Error | z value | Pr(>\|z\|) |
| --- | --- | --- | --- | --- |
| (Intercept) | -2.45122 | 0.84472 | -2.902 | 0.003710 |
| Cholearly | -1.57245 | 1.17473 | -1.339 | 0.180713 |
| Chollate | -0.16256 | 1.17275 | -0.139 | 0.889757 |
| Day | 0.28672 | 0.07755 | 3.697 | 0.000218 |
| Cholearly:Day | -0.29539 | 0.10864 | -2.719 | 0.006551 |
| Chollate:Day | -0.22960 | 0.10959 | -2.095 | 0.036159 |

**Effect sizes:**

The ‘early’ cholestyramine treatement reduced the normalized mean proportion of resistant *E. faecium* relative to the no cholestyramine control by 0.14 (95% CI ± 0.13) at Day 2, 0.30 (± 0.17) at Day 4, 0.43 (± 0.14) at Day 6, 0.32 (± 0.19) at Day 8, and 0.51 (± 0.21) at Day 14., The ‘late’ cholestyramine reduced the normalized mean proportion resistant *E. faecium* compared to the no cholestyramine control by 0.06 (95% CI ± 0.17) at Day 2, 0.08 (± 0.26) at Day 4, -0.05 (± 0.287) at Day 6, 0.02 (± 0.30) at Day 8, and 0.32 (± 0.21) at Day 14.

**Model 2: Effect of cholestyramine timing on shedding of daptomycin-resistant *E. faecium* in daptomycin-treated mice**

**Input data:** Fig3_CFUdata_choltiming.csv

**Data preparation:**

1. For each mouse, calculate AUC for resistant bacteria (AUC_R)

AUC(x=data$Day, y=data$resist.per10mg)

**Model Structure in R:**

glm(log10(AUC_R) ~ Chol, data = data[data$Antibiotic=="daptomycin",])

summary()

|  | Estimate | Std. Error | t-value | p-value |
| --- | --- | --- | --- | --- |
| (Intercept) | 7.6760 | 0.2847 | 26.961 | < 2e-16 |
| Cholearly | -1.1178 | 0.4026 | -2.776 | 0.00987 |
| Chollate | -0.6473 | 0.4026 | -1.608 | 0.11955 |

**Model 3: Effect of cholestyramine timing on total shedding (S+R) of *E. faecium* in daptomycin-treated mice**

**Input data:** Fig3_CFUdata_choltiming.csv

**Data preparation:**

1. For each mouse, calculate AUC for total *E. faecium* (AUC_T)

AUC(x=data$Day, y=data$total.per10mg)

**Model Structure in R:**

glm(log10(AUC_T) ~ Chol, data = data[data$Antibiotic=="daptomycin",])

summary()

|  | Estimate | Std. Error | t-value | p-value |
| --- | --- | --- | --- | --- |
| (Intercept) | 8.03003 | 0.14262 | 56.304 | < 2e-16 |
| Cholearly | -0.03546 | 0.20169 | -0.176 | 0.862 |
| Chollate | -0.02951 | 0.20169 | -0.146 | 0.885 |

**Model 4: Effect of cholestyramine timing on total shedding (S+R) of *E. faecium* in mice *without* daptomycin treatment**

**Input data:** Fig3_CFUdata_choltiming.csv

**Data preparation:**

1. For each mouse, calculate AUC for total *E. faecium* (AUC_T)

AUC(x=data$Day, y=data$total.per10mg)

**Model structure in R:**

glm(log10(AUC_T) ~ Chol, data = data[data$Antibiotic=="control",])

summary()

|  | Estimate | Std. Error | t-value | p-value |
| --- | --- | --- | --- | --- |
| (Intercept) | 8.1833 | 0.2152 | 38.025 | < 2e-16 |
| Cholearly | -0.4066 | 0.3043 | -1.336 | 0.193 |
| Chollate | -0.3827 | 0.3043 | -1.257 | 0.219 |
